# Expression of Filaments of the *Geobacter* Extracellular Cytochrome OmcS in *Shewanella oneidensis*

**DOI:** 10.1101/2023.10.26.564108

**Authors:** Tong Lin, Wenqi Ding, Danni Zhang, Yun Yang, Feng Li, Dake Xu, Derek R. Lovley, Hao Song

## Abstract

The physiological role of *Geobacter sulfurreducens* extracellular cytochrome filaments is a matter of debate and the development of proposed electronic device applications of cytochrome filaments awaits methods for large-scale cytochrome nanowire production. Functional studies in *Geobacter sulfurreducens* are stymied by the broad diversity of redox-active proteins on the outer cell surface and the redundancy and plasticity of extracellular electron transport routes. *G. sulfurreducens* is a poor chassis for producing cytochrome nanowires for electronics because of its slow, anaerobic growth. Here we report that filaments of the *G. sulfurreducens* cytochrome OmcS can be heterologously expressed in *Shewanella oneidensis*. Multiple lines of evidence demonstrated that a strain of *S. oneidensis*, expressing the *G. sulfurreducens* OmcS gene on a plasmid, localized OmcS on the outer cell surface. Atomic force microscopy revealed filaments with the unique morphology of OmcS filaments emanating from cells. Electron transfer to OmcS appeared to require a functional outer-membrane porin-cytochrome conduit. The results suggest that *S. oneidensis*, which grows rapidly to high culture densities under aerobic conditions, may be a suitable for development of a chassis for producing cytochrome nanowires for electronics applications and may also be good model microbe for elucidating cytochrome filament function in anaerobic extracellular electron transfer.

---

Extracellular filaments comprised of multi-heme *c*-type cytochromes are of interest because of their potential role in electron transfer to insoluble electron acceptors and because they could have applications as sustainably produced conductive nanowires in electronic devices (Atkinson, Chavez, Ninman, & El-Naggar, 2023; Gralnick & Bond, 2023; Lovley & Holmes, 2022; Lovley & Yao, 2021).

Cytochrome filaments have been studied most in *Geobacter sulfurreducens*, which has served as a model microbe to elucidate the physiology and ecology of the electroactive *Geobacter* species that play an important role the biogeochemistry of diverse anaerobic environments and bioelectrochemical systems harvesting electricity from waste organic matter (Lovley & Holmes, 2022).

Three *G. sulfurreducens* multi-heme *c*-type cytochrome filaments are known to assemble into filaments OmcS (Filman et al., 2019; Wang et al., 2019), OmcZ (Wang, Chan, et al., 2022; Yalcin et al., 2020), and OmcE (Wang, Mustafa, et al., 2022). It has been suggested that these three cytochrome filaments are conduits for long-range electron transport to electron acceptors at distance from the cell (Wang, Chan, et al., 2022; Wang et al., 2019; Wang, Mustafa, et al., 2022; Yalcin et al., 2020). However, analysis of functional genetic studies have challenged this concept, summarizing the experimental evidence that refutes the concept that OmcS, OmcZ, or OmcE filaments account for the long-range conductivity in current producing biofilms (Lovley & Holmes, 2022). Furthermore, none of the filament-forming cytochromes are essential for the reduction of Fe(III) oxide, one of the most environmentally significant extracellular electron acceptors in the native soil and sediment habitats of *Geobacter* species (Liu et al., 2022; Schwarz et al., 2023).

The abundance of diverse *G. sulfurreducens* outer-surface redox-active proteins, unexpected pleiotropic impacts of gene deletions on the expression or localization of other electron transport proteins, and the likelihood of compensatory mutations to adapt to gene deletions can confound functional genetic studies (Lovley & Holmes, 2022). Thus, there is a strong need for ‘bottom up’ approaches in which the function and interaction of *G. sulfurreducens* outer-surface proteins are studied in a more controlled manner in a heterologous host (Lovley & Holmes, 2022).

Expression of cytochrome filaments in a heterologous host will also be necessary in order to explore potential applications of cytochrome filaments in electronic devices. Cytochrome filament production with *G. sulfurreducens* is impractical because of its slow anaerobic growth and the contamination of cytochrome filament preparations with electrically conductive pili which are nearly 10-fold more abundant in rapidly growing *G. sulfurrreducens* (Liu, Walker, Nonnenmann, Sun, & Lovley, 2021; Tan et al., 2016). *G. sulfurreducens* gene deletion mutants that do not express pili also do not express OmcS filaments. OmcS was previously expressed in *Synechococcus elongatus* (Dong, Lee, Gaffney, Liou, & Minteer, 2021; Sekar, Jain, Yan, & Ramasamy, 2016), but OmcS filaments were not observed.

Therefore, the possibility of expressing *G. sulfurreducens* OmcS filaments in a heterologous host was evaluated with *Shewanella oneidensis*, which expresses an abundance of native multi-heme *c*-type cytochromes, but not its own extracellular cytochrome filaments (Gralnick & Bond, 2023; Lovley & Holmes, 2022). The *G*.

*sulfurreducens* gene for OmcS was adapted for *S. oneidensis* codon usage and expressed in *S. oneidensis* (Figure 1A) on a plasmid under control of the inducer IPTG (see Supporting Information for detailed experimental methods). This strain was designated strain OmcS (Table S1, supporting material). Analysis of protein extracts from cells grown aerobically in LB medium with SDS-PAGE gel separation and silver staining (Figure 1B) revealed the expression of a protein with an apparent molecular weight of ∼50 kDa, the expected size of OmcS (Mehta, Coppi, Childers, & Lovley, 2005; Qian et al., 2011). There was no equivalent band in the control strain MY, carrying the empty plasmid without the OmcS gene (Figure 1B). Liquid chromatography coupled with tandem mass spectrometry (LC-MS/MS) of the peptides collected from digested protein extracts detected four unique OmcS peptides representing 21.7% coverage (94/432 amino acids) (Figure S1), further confirming OmcS expression. Heme-staining of proteins separated on gels revealed a heme-containing protein at ∼50 kDa for strain OmcS, but not strain MY (Figure 1C). When cells were treated with proteinase K in a manner previously demonstrated to selectively remove outer surface proteins (Qian, Reguera, Mester, & Lovley, 2007) the ∼50 kDa band decreased with increasing treatment time (Figure 1D). In contrast, the 62 kDa periplasmic *c*-type cytochrome FccA (Delgado, Paquete, Sturm, & Gescher, 2019) remained intact throughout the proteinase K treatment (Figure 1D), indicating that the outer membrane was not disrupted during proteolysis. These results demonstrated that OmcS was exposed on the outer surface of *S. oneidensis*, which was consistent with OmcS localization in *G. sulfurreducens* (Mehta et al., 2005).

**Figure 1.**
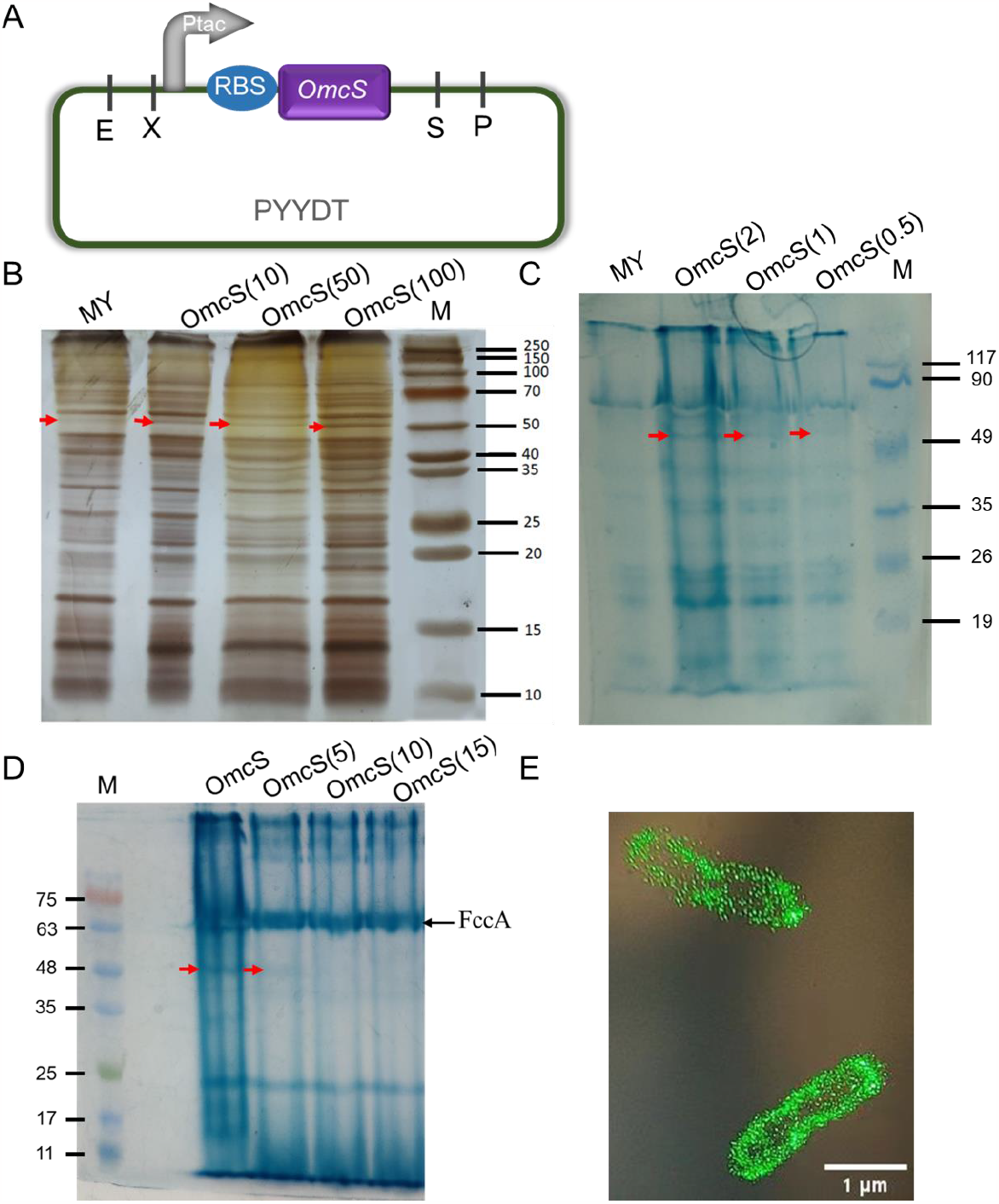
Expression of the *Geobacter sulfurreducens* multi-heme *c*-type cytochrome OmcS in *Shewanella oneidensis* strain OmcS. (A) Plasmid map of the PYYDT vector expressing the *G. sulfurreducens* OmcS gene under the control of the inducible promoter P_tac_ in strain OmcS. E, X, S, P designate the restriction enzyme sites of EcoRI, XbaI, SpeI, SbfI /SdaI, respectively. (B) Total protein separated by SDS-PAGE gel and then silver stained. OmcS (10), OmcS (50), OmcS (100) designate the OmcS protein expressed in the recombinant *S. oneidensis* OmcS strain with IPTG inducer concentrations of 10 μM, 50 μM and 100 μM, respectively. Arrows designate the expected position of the OmcS protein band. (C) Heme staining of separated cell lysate proteins with arrows indicating the expected position of OmcS. OmcS (2), OmcS (1), OmcS (0.5) designate that the culture OD_600_ of the OmcS strain with 10 μM IPTG induction, was 2.0, 1.0 or 0.5, respectively. (D) Heme-stained protein profile of whole cells treated with proteinase K. Lane designations are: OmcS, untreated control of strain OmcS; OmcS (5), OmcS (10), OmcS (15), strain OmcS treated with proteinase K (1 U/mL) for 5, 10, and 15 min, respectively. (E) Confocal laser scanning micrograph of the recombinant *S. oneidensis* strain in which OmcS was fused with the super folder green fluorescent protein (sfGFP) (i.e., *S. oneidensis* MR-1 harboring the plasmid P_tac_-*OmcS-sfGFP*-PYYDT).

A strain with the OmcS gene fused with a C-terminal super fluorescent protein GFP (*OmcS-sfGFP*) displayed substantial fluorescence in the outer cell surface (Figure 1E), further suggesting an outer-surface OmcS localization. Fluorescence at distance from the cells, which might be indicative of OmcS filaments was not observed, but this result might have reflected a lack of sufficient microscope resolution or the addition of the fluorescent protein interfering with filament assembly.

Therefore, the potential for strain OmcS to express OmcS filaments was further evaluated with a high-resolution atomic force microscopy method that was previously successful in identifying OmcS filaments emanating from *G. sulfurreducens* (Liu et al., 2021). OmcS filaments have a diameter of ca. 4 nm and a characteristic 20 nm longitudinal pitch (Filman et al., 2019; Liu et al., 2021; Wang et al., 2019). Wild-type *S. oneidensis* produces type IV and Msh (mannose sensitive hemagglutinin) pili as well as flagella (Bouhenni et al., 2010). Atomic force microscopy revealed numerous putative pili and flagella emanating from both strain OmcS (Figure 2, Figures S2, S3) and the control MY strain (Figure S4). These filaments were readily distinguished from the OmcS filaments based on filament height (i.e., diameter) and the lack of the characteristic 20 nm longitudinal pitch (Figure 2, Figures S2, S3, S4). About 10% of the filaments emanating from strain OmcS had the unique morphology (4 nm diameter coupled with 20 nm longitudinal pitch) of OmcS filaments (Figure 2, Figures S2, S3). This is similar to the percentage of OmcS filaments emanating from *G. sulfurreducens* (Liu et al., 2021). These results demonstrated that not only did strain OmcS express OmcS, but also that at least some of the OmcS was assembled into filaments in a manner similar to that observed for *G. sulfurreducens*.

**Figure 2.**
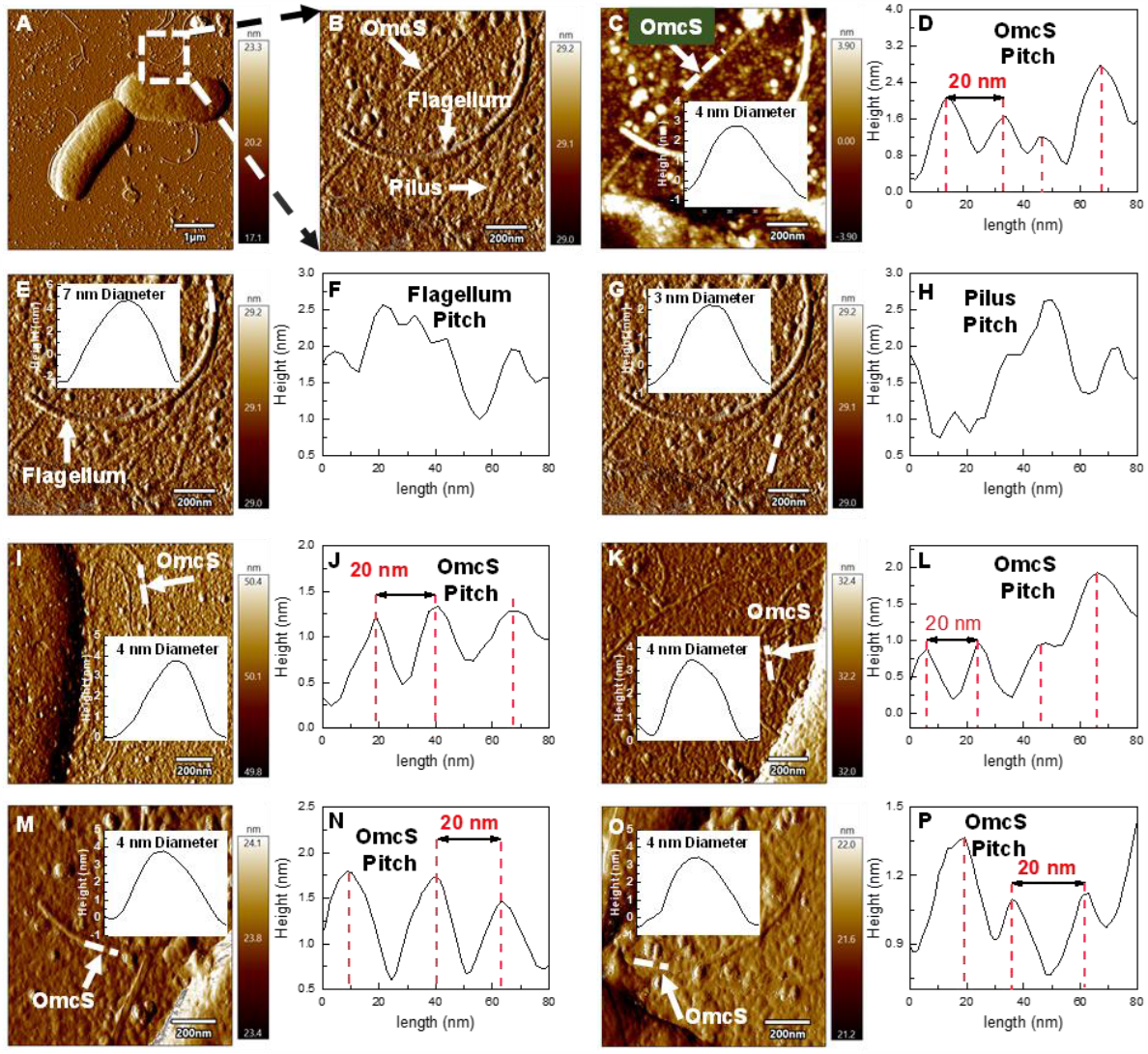
Atomic force microscopy analysis of images emanating from the *S. oneidensis* strain OmcS. (A) Cells with diverse filaments. (B) Identification of putative filament types. (C) Higher magnification and height measurement (inset) of putative OmcS filament. (D) Profile along the length of the OmcS filament (taken where designated with dashed white line in panel C demonstrating the 20 nm pitch characteristic of OmcS filaments. (E) Higher magnification and height measurement (inset) of larger filament shown in panels A and B, designated here as a flagellum to distinguish it from thinner filaments. (F) Profile along the length of putative flagellum (taken where designated with dashed white line in panel E). (G) Higher magnification and height measurement (inset) of one of the numerous thinner filaments emanating from cells in panels A and B. (H) Profile along the length of 3 nm filament (taken where designated with dashed line in panel G. (I-P) Additional images of OmcS filaments emanating from cells displaying 4 nm diameter and 20 nm pitch along the filament length.

In order to determine whether expression of OmcS impacted on extracellular electron transfer, cell suspensions introduced into a bioelectrochemical system with lactate as the electron donor and the anode as a sole electron acceptor. Strain OmcS produced 2-fold more current than the control strain (Figure 3 A). Viable cell counts, and protein analysis (Figure 3B), demonstrated that the number of strain OmcS or strain MY cells attached to anodes were not significantly different (*p* > 0.05), as was also apparent from scanning electron microscopy (Figures 3 C and D). Thus, the per cell power output of strain OmcS was higher than for strain MY. For example, based on the viable cell count (Fig. 3B), the power output of by a single OmcS cell was (4.723 ± 0.21) × 10^−10^ W/m^2^, 2.336-fold higher than that of a single cell of strain MY ((2.021 ± 0.13) × 10^−10^ W/m^2^).

**Figure 3.**
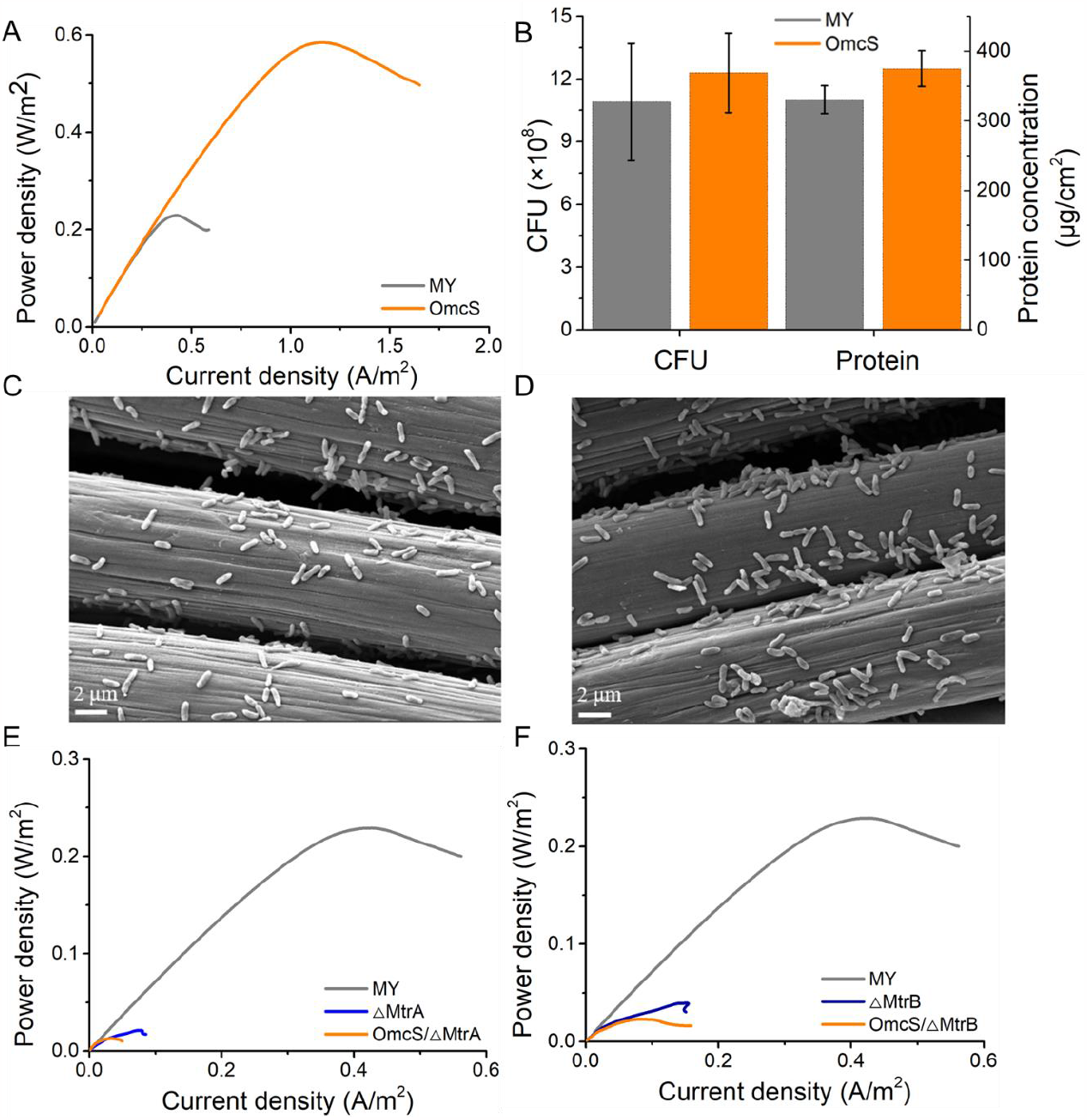
Growth and activity of *S. oneidensis* strain OmcS and the control MY strain on electrodes. (A) Strain OmcS and strain MY current generation. (B) Colony forming units (CFU) and protein recovered from biofilms of strains OmcS and MY (C) Scanning electron micrograph of strain MY anode biofilm. (D) Scanning electron micrograph of strain OmcS anode biofilm. (E) Current production with strain ΔMtrA in which the gene for MtrA, the periplasmic-facing cytochrome component of the porin-cytocytochrome conduit, was deleted, and strain OmcS/ΔMtrA in which the gene for OmcS was expressed in strain ΔMtrA. (F) Current production with strain ΔMtrB in which the gene for MtrB, the porin protein of the porin-cytochrome conduit was deleted, and strain OmcS/ΔMtrB in which the OmcS gene was expressed in strain ΔMtrB.

Scanning electron microscopy also revealed that current-producing cells of both strain MY and strain OmcS were in direct contact with the anode (Figures 3 C and D). This observation is consistent with previous studies with *G. sulfurreducens* that have suggested that OmcS can enhance short-range electron exchange between cells and anodes when current production is low and biofilms are thin (Holmes et al., 2006), but that OmcS is not involved in long-range electron transport through thick, electrically conductive biofilms (Malvankar, Tuominen, & Lovley, 2012; Nevin et al., 2009).

Both *S. oneidensis* and *G. sulfurreducens* have porin-cytochrome complexes for transferring electrons across the outer membrane (Gralnick & Bond, 2023; Lovley & Holmes, 2022). Previous studies (Bretschger et al., 2007) demonstrated that deletion of the gene for MtrA, the periplasmic-facing multi-heme cytochrome of the *S. oneidensis* strains porin-cytochrome complex, or MtrB, the porin protein, inhibited electron transfer through the *S. oneidensis* porin-cytochrome conduit, limiting current production. These same gene deletions also limited current production in our system (Figures 3E and F). Current production remained limited when OmcS was expressed in the MtrA- or MtrB-deficient strains (Figures 3E and F). This result is consistent with the concept that cytochrome filaments or conductive pili do not transport electrons across the outer membrane, but rather accept electrons transported to the outer surface via porin-cytochrome conduits (Gralnick & Bond, 2023; Lovley & Holmes, 2022).

The finding that *G. sulfurreducens* OmcS filaments can be heterologously expressed in *S. oneidensis* simply by introducing this gene on a plasmid suggests that it will be possible to construct strains for OmcS filament production for electronics applications, and to study OmcS filament function in a microbe with a less complex array of outer-surface redox proteins than what must be dealt with in *G. sulfurreducens*. Optimizing OmcS production will require substantial additional genetic modifications to eliminate the production of other *S. oneidensis* filaments, such as pili and flagella. These filaments have previously been removed in other *S. oneidensis* strain constructions (Bouhenni et al., 2010; Szmuc et al., 2023). Further optimization might include incorporating the OmcS gene within the chromosome under the control of a strong native promoter and deletion of genes for other unnecessary cellular components to free up biosynthetic resources for OmcS filament production. It seems likely that the other *G. sulfurreducens* cytochrome filaments, OmcZ and OmcE, might be produced in a similar manner.

The successful expression of OmcS filaments in *S. oneidensis*, coupled with the recent demonstration that *G. sulfurreduens* conductive pili can also be heterologously expressed in *S. oneidensis* (Szmuc et al., 2023), suggests that *S. oneidensis* could be an excellent foundational strain for the previously proposed (Lovley & Holmes, 2022) ‘bottom up’ approach of evaluating the function and interactions of multiple *G. sulfurreducens* electron transfer components in a microbe other than *G. sulfurreducens*. Conducting such studies in *S. oneidensis*, which is easier to grow and genetically manipulate than *G. sulfurreducens*, could greatly accelerate developing an understanding of how *G. sulfurreducens* is so effective in extracellular electron exchange (Lovley & Holmes, 2022).

## ACKNOWLEDGMENTS

This work was supported by the National Key Research and Development Program of China (2018YFA0901300), the National Natural Science Foundation of China (NSFC 22378305, 32071411 and 21621004), the Natural Science Foundation of Hebei Province (B2020408005), and the Science and Technology Project of Hebei Education Department (QN2021125).

## AUTHOR CONTRIBUTIONS

T. L., W. D., and D. Z. contributed equally to this work. T. L. and W. D. designed the project, performed experiments, analyzed data, and drafted the manuscript. D. Z. conducted the atomic force microscopy studies. Y. Y., D. Z., F. L. and D. X. helped for performing several experiments. D. R. L. and H. S. designed and supervised the project, analyzed data, and revised the manuscript.

## ETHICS STATEMENT

This study did not use animals.

## DATA AVAILABILITY

All data are available within the article or supplementary information file(s). All other data supporting the findings of this study are available from the corresponding authors upon reasonable request.

## References

Atkinson, J. T., Chavez, M. S., Ninman, C. M., & El-Naggar, M. Y. (2023). Living electronics: A catalogue of engineered living electronic components. Microb Biotechnol, 16, 507–533.

Bouhenni, R. A., Vora, G. J., Biffinger, J. C., Shirodkar, S., Brockman, K., Ray, R., … Saffarini, D. A. (2010). The role of Shewanella oneidensis MR-1 outer surface structures in extracellular electron transfer. Electroanalysis, 22, 856–864.

Bretschger, O., Obraztsova, A., Sturm, C. A., Chang, I. S., Gorby, Y. A., Reed, S. B., … Nealson, K. H. (2007). Current production and metal oxide reduction by Shewanella oneidensis MR-1 wild type and mutants. Appl. Environ. Microbiol., 73, 7003–7012.

Delgado, V. P., Paquete, C. M., Sturm, G., & Gescher, J. (2019). Improvement of the electron transfer rate in Shewanella oneidensis MR-1 using a tailored periplasmic protein composition. Bioelctrochemistry, 129, 18–25.

Dong, F., Lee, Y. S., Gaffney, E. M., Liou, W., & Minteer, S. D. (2021). Engineering cyanobacterium with transmembrane electron transfer ability for bioelectrochemical nitrogen fixation. ACS Catalysis, 11, 13169–13179.

Filman, D. J., Marino, S. F., Ward, J. E., Yang, L., Mester, Z., Bullitt, E., … Strauss, M. (2019). Cryo-EM reveals the structural basis of long-range electron transport in a cytochrome-based bacterial nanowire. Communications Biology, 2, 219.

Gralnick, J. A., & Bond, D. R. (2023). Electron transfer beyond the outer membrane: putting electrons to rest. Annual Review of Microbiology, 77, 517–539.

Holmes, D. E., Chaudhuri, S. K., Nevin, K. P., Mehta, T., Methe, B. A., Liu, A., … Lovley, D. R. (2006). Microarray and genetic analysis of electron transfer to electrodes in Geobacter sulfurreducens. Env. Microbiol., 8(10), 1805–1815.

Liu, X., Walker, D. J. F., Li, Y., Meier, D., Pinches, S., Holmes, D. E., & Smith, J. A. (2022). Cytochrome OmcS is not essential for extracellular electron transport via conductive pili in Geobacter sulfurreducens strain KN400. Appl Environ Microbiol, 88, e01622–21.

Liu, X., Walker, D. J. F., Nonnenmann, S., Sun, D., & Lovley, D. R. (2021). Direct observation of electrically conductive pili emanating from Geobacter sulfurreducens. mBio, 12, e02209–21.

Lovley, D. R., & Holmes, D. E. (2022). Electromicrobiology: The ecophysiology of phylogenetically diverse electroactive microorganisms. Nature Reviews Microbiology, 20, 5–19.

Lovley, D. R., & Yao, J. (2021). Intrinsically conductive microbial nanowires for ‘green’ electronics with novel functions. Trends in Biotechnology, 39, 940–952.

Malvankar, N. S., Tuominen, M. T., & Lovley, D. R. (2012). Lack of involvement of c-type cytochromes in long-range electron transport through conductive biofilms and nanowires. Energy. Environ. Sci., 5, 8651 –8659.

Mehta, T., Coppi, M. V., Childers, S. E., & Lovley, D. R. (2005). Outer membrane ctype cytochromes required for Fe(III) and Mn(IV) oxide reduction in Geobacter sulfurreducens. Appl. Environ. Microbiol., 71, 8634–8641.

Nevin, K. P., Kim, B.-C., Glaven, R. H., Johnson, J. P., Woodard, T. L., Methé, B. A., … Lovley, D. R. (2009). Anode biofilm transcriptomics reveals outer surface components essential for high current power production in Geobacter sulfurreducens fuel cells. PLoS One, 4, e5628.

Qian, X., Mester, T., Morgado, L., Arakawa, T., Sharma, M. L., Inoue, K., … Lovley, D. R. (2011). Biochemical characterization of purified OmcS, a C-type cytochrome required for insoluble Fe(III) reduction in Geobacter sulfurreducens. Biochim. Biophys. Acta, 1807, 404–412.

Qian, X., Reguera, G., Mester, T., & Lovley, D. R. (2007). Evidence that OmcB and OmpB of Geobacter sufurreducens are outer membrane surface proteins. FEMS Microb Lett, 277, 21–27.

Schwarz, I. A., Alsaqri, B., Lekbach, Y., Henry, K., Gorman, S., Woodard, T. L., … Lovley, D. R. (2023). Lack of physiological evidence for cytochrome filaments functioning as conduits for extracellular electron transfer. manuscript submitted.

Sekar, N., Jain, R., Yan, Y., & Ramasamy, R. P. (2016). Enhanced photobioelectrochemical energy conversion by genetically engineered cyanobacteria. Biotechnology and Bioengineering, 113, 675–679.

Szmuc, E., Walker, D. J. F., Kireev, D., Akinwande, D., Lovley, D. R., Keitz, B. K., & Ellington, A. (2023). Engineering Geobacter pili to produce metal:organic filaments. Biosensors and Bioelectronics, 222, 114993.

Tan, Y., Adhikari, R. Y., Malvankar, N. S., Ward, J. E., Nevin, K. P., Woodard, T. L., … Lovley, D. R. (2016). The low conductivity of Geobacter uraniireducens pili suggests a diversity of extracellular electron transfer mechanisms in the genus Geobacter. Front Microbiol, 7, 980.

Wang, F., Chan, C. H., Suciu, V., Mustafa, K., Ammend, M., Si, D., … Bond, D. R. (2022). Structure of Geobacter OmcZ filaments suggests extracellular cytochrome polymers evolved independently multiple times. eLife, 11, e8155.

Wang, F., Gu, Y., O’Brien, J. P., Yi, S. M., Yalcin, S. E., Srikanth, V., … Malvankar, N. S. (2019). Structure of microbial nanowires reveals stacked hemes that transport electrons over micrometers. Cell, 177, 361–369.

Wang, F., Mustafa, K., Suciu, V., Joshi, K., Chan, C. H., Choi, S., … Bond, D. R. (2022). Cryo-EM structure of an extracellular Geobacter OmcE cytochrome filament reveals tetrahaem packing. Nature Microbiology, 7, 1291–1300.

Yalcin, S. E., O’Brien, J. P., Gu, Y., Reiss, K., Yi, S. M., Jain, R., … Malvankar, N. S. (2020). Electric field stimulates production of highly conductive microbial OmcZ nanowires. Nature Chemical Biology, 16, 1136–1142.

