## Supplemental Information for "Expression of Filaments of the *Geobacter* Extracellular Cytochrome OmcS in *Shewanella oneidensis*"

#### Experimental Details

##### In vitro gene synthesis, plasmid construction and transformation

The *OmcS* gene of *G. sulfurreducens* extracted from KEGG (Kanehisa & Goto, 2000) was input into JCat ((Java Codon Adaptation Tool) (Grote et al., 2005) to adapt codon usage for expression in *S. onedensis*, avoiding the restriction enzyme sites of EcoRI, XbaI, SpeI, and SdaI (SbfI). The sequence was synthesized in vitro as a BioBrick gene, which included an upstream prefix EcoRI and XbaI, the ribosome binding site (RBS, BBa_B0034, iGEM), and a downstream suffix SpeI and SdaI. The *OmcS* gene fused with a C-terminal super fluorescent protein GFP (*OmcS-sfGFP*) was also synthesized *in vitro* using a similar BioBrick method.

The synthesized BioBrick genes were inserted into the previously described vector PYYDT (Yang et al., 2015). Plasmids were transformed into *E. coli* WM3064, the dap auxotroph strain (Newman & Saltikov, 2003), which was grown with 100 μg/mL 2, 6-diaminopimelic acid, then transferred into *S. onedensis* by conjugation. The following previously described (Bretschger et al., 2007) *S. oneidensis* strains were also studied:ΔMtrA (SO1777), ΔMtrB (SO1776), as well as *S. oneidensis* strain MY (49). Unless otherwise specified, strains were grown in LB medium at 30°C with shaking at 200 rpm.

**Bioelectrochemical studies**

For current production studies, cells were grown in a two-chamber (140 ml chamber volumes) microbial fuel cell in which a carbon cloth anode (1.0 cm × 1.0 cm) was separated from a carbon cloth cathode (2.5 cm × 3.0 cm) with a Nafion 117 membrane. Prior to inoculation the system was treated with 3% hydrogen peroxide at 80°C for 1 h then washed with distilled water, and 0.5 M sulfuric acid and then washed with sterile distilled water. The cathodic electrolyte was comprised of 50 mM $K_3[Fe(CN)_6]$, 50 mM $KH_2PO_4$, and 50 mM $K_2HPO_4$. The anode and cathode were connected with a 2 kΩ resistor. The microbial fuel cells were incubated at 30°C. Output voltages were continuously recorded with MPS110001data acquisition cards.

To initiate current production studies single colonies of *S. oneidensis* were grown in LB medium overnight. Then a 1% inoculum was transferred into fresh LB broth containing 50 μg/mL kanamycin and ITPG inducer (10 μM). After 12 h, the cell suspension concentration was adjusted to $OD_{600}$ = 0.8, and dispersed into the anode chamber containing mineral-vitamin mix (Baron, LaBelle, Coursolle, Gralnick, & Bond, 2009), M9 buffer, 18 mM lactate, 50 μg/mL kanamycin and IPTG inducer (10 μM). The anode chambers were purged with $N_2$ gas to remove oxygen.

Cyclic voltammetry with an Ag/AgCl (KCl saturated) reference electrode was conducted in a three-electrode configuration. Linear sweep voltammetry, in which the potential decreased from the open circuit potential (OCP) to −0.3 V at 0.1 mV/s, was applied to obtain the polarization curves with a CHI1000C electrochemical workstation (CH Instrument, Shanghai, China).

Electron uptake from cathodes was evaluated with a three-electrode electrochemical cell continuously stirred with a multipoint magnetic stirrer (Xie et al., 2020). The working electrode of the electrochemical cell was a biofilm grown in current-producing mode on carbon cloth. The counter electrode was a carbon cloth in the electrolyte, which was comprised of 50 mM $KH_2PO_4$, 50 mM $K_4[Fe(CN)_6]$ and 50 mM $K_2HPO_4$. The Ag/AgCl (KCl saturated) was the reference electrode. The working electrode was poised at −0.56 V (*vs.* Ag/AgCl), and the inward current was measured with a CHI1000C electrochemical workstation.

**Colony forming units (CFU) counting and bicinchoninic acid (BCA) measurement of total proteins on anodes**

To estimate the number of viable cells growing on biofilms, anodes were placed in 5 mL of 0.9% sterile NaCl solution and vortexed for 2 min. Serial dilutions were spread on LB agar plates that contained 50 μg/mL kanamycin. *S. oneidensis* colonies forming on the plates were counted after incubation at 30°C for 12-24 h. For protein determinations, anodes were placed in 5 mL of 0.2 M NaOH at 96℃ for 20 min to lyse cells. After cooling to room temperature, the protein in the lysate was determined with the BCA protein assay kit (Novagen, China).

#### Gel electrophoresis and protein identification

Cells were collected with centrifugation (12,000 rpm, 10 min, 4°C). The pellet was frozen with liquid nitrogen and ground into a powder, which was lysed on ice for 20 min in 1 mL Radio immunoprecipitation assay (RIPA) lysis buffer, which was comprised of 1 mM phosphatase inhibitor mixture, 1 mM protease inhibitor mixture, 1 mM phenylmethanesulfonyl fluoride and 1 mM dithiothreitol (DTT). The lysate was then treated with ultrasound (CV 334, Scientz, China) for 5 min at 20% power, and centrifuged at 10,000 rpm for 15 min at 4 ℃. Proteins in the supernatant were separated with sodium dodecyl sulfate polyacrylamide gel electrophoresis (SDS-PAGE) with 12% gels using a vertical electrophoresis system at a constant voltage of 90 V for 3 h. The gels were stained with the silver staining method (Heukeshoven & Dernick, 1985).

Proteins in the cell lysate were identified with slight modifications to previously described methods (Foster & Mann, 2006; Gundry et al., 2010; Ludwig, Schroll, & Hummon, 2018). Protein was concentrated with a 10 kD ultrafiltration tube (Milipore, USA), reduced with DTT (final concentration 25 mM) at 37°C for 1 h. Proteins were then alkylated with iodoacetamide (final concentration 50 mM IAA) and the sample was vortexed at 25°C for 30 min in the dark. The sample was washed four times with 200 mM triethylammonium bicarbonate buffer (TEAB), centrifuging the sample at 12,000 rpm for 10 min and discarding the flow-through at each step. Proteins were digested trypsin overnight at 37°C. Digested peptides were eluted with TEAB and centrifugation (12,000 rpm for 10 min), a process that was repeated 3 times. Digested peptides were stored at −20°C. Samples were desalted, with a $C_{18}$ solid-phase extraction column (Milipore, USA) as previously described (Foster & Mann, 2006) and the desalted peptide sample was vacuum freeze-dried. Peptides components were dissolved in 0.1% formic acid solution and separated with a low pH reversed-phase $C_{18}$ capillary chromatography (75 μm × 200 mm, 3 μm). The peptides were separated with a gradient elution employing solvent A (99.9% water and 0.1% formic acid) and solvent B (80% acetonitrile, 19.9% water and 0.1% formic acid) with the solvent A and solvent B with ratios shifted over a 90 min run from 93:7 to 5:95 v/v. The mass spectrometry (Orbitrap Q-Exactive, Thermo, USA) was performed using data-dependent acquisition MS/MS scans with a mass range of 300–1800 m/z (Aebersold & Mann, 2003; Gstaiger & Aebersold, 2009). Peptide masses were queried against entries in the uniport *Geobacter sulfurreducens* PCA database with MaxQuant.

For analysis of heme-containing proteins, the *S. oneidensis* strain OmcS in LB medium was induced with 10 μM IPTG for 12 h. Cell were collected with centrifugation (4000 rpm, 20 min, 4°C), washed three times with PBS buffer, 5 × Native Gel Sample Loading Buffer (Beyotime) was added to the samples and the $OD_{600}$ was adjusted to 2.0, 1.0, or 0.5. Samples (20 μL) were separated with SDS-PAGE in 12% gel (SDS-PAGE Gel Quick Preparation Kit, Beyotime). Heme-containing proteins were visualized as previously described (Thomas, Ryan, & Levin, 1976). Gels were incubated in a solution of 0.03 g tetramethylbenzidine dissolved in 15 mL methanol mixed with 35 mL of 0.5 M sodium acetate in the dark for 2 h. $H_2O_2$ (300 μL) was added and heme-containing bands observed after 1 h.

In order to determine which heme proteins might be exposed on the outer cell surface and thus susceptible to protease treatment (Qian, Reguera, Mester, & Lovley, 2007), strain OmcS was grown in LB medium with IPTG (10 µM) for 12 h. Cells were collected by centrifugation (4000 rpm, 20 min, 4℃) washed twice with PBS buffer, and resuspended in 10 mM HEPES (pH 7.2-7.4) containing 500 µM $MgCl_2$. The strains were incubated with or without 1 U/mL proteinase K at 4℃, for different lengths of time (i.e., 5, 10 and 15 min). A protease inhibitor was then added to stop the proteolytic reaction. Cells were collected with centrifugation (4000 rpm, 10 min, 4°C), washed twice, and resuspended in HEPES buffer. Proteins separated with SDS-PAGE and heme proteins visualized with heme staining as described above.

**Microscopy**

To evaluate OmcS localization, the recombinant *S. oneidensis* containing a gene encoding a protein in which OmcS was fused with the super folder GFP protein was induced to express the gene for 16 h. Cells were collected by centrifugation (5000 rpm, 10 min, 4℃), the pellet was washed three times with 0.9% sterile NaCl. Cells were then resuspended in 0.9% NaCl. Images at laser wavelengths of 485 nm and 525 nm were captured with a Nikon A1R+ confocal laser scanning microscope and processed with NIS-Elements software.

To directly visualize filaments emanating from cells, strain OmcS or strain MY were cultured in LB broth with 50 µg/mL kanamycin and 0.01 mM IPTG for 8-12 h. Cells were collected by centrifugation (3050 *g*, 10 min, 4℃) and washed three times with PBS buffer. As previously described (Liu, Walker, Nonnenmann, Sun, & Lovley, 2021), a 45 µL aliquot of culture was drop cast onto silicon wafers that were coated with a 35 nm layer of platinum. After 15 min, excess liquid was removed with a pipette and the sample was washed with 1 mL of deionized water. Excess water was absorbed with filter paper and the preparation was allowed to air dry. The filaments were observed with tapping mode (AC-air topography) under repulsive force with a Pt/Ir-coated tip (PtSi-FM, NanoWorld AG) at a ~1.9 N/m spring force constant and ~73 kHz resonance frequency with Cypher ES, atomic force microscope (Asylum Research, Oxford Instrument).

To visualize anode biofilms, small pieces of carbon cloth anodes were fixed with 2.5% glutaraldehyde for 1h, dehydrated in series of ethanol solutions (25%, 50%, 75%, 95% and 100%), and then vacuum freeze dried. The samples were coated with Au before imaging with a TESCAN MIRA scanning electron microscope at 5 kV accelerating voltage (EHT), 4.88 mm working distance (WD) and In-Beam SE detector.

**Table S1**. The recombinant strains and their genotypes used in this study

| **Strain** | **Relevant characteristic(s)** | **Source** |
|---|---|---|
| MY | *S. oneidensis* MR-1 harbouring the empty plasmid PYYDT | Lab stock |
| OmcS | *S. oneidensis* MR-1 harbouring the plasmid $P_{tac}$-*OmcS*-PYYDT | This work |
| ΔMtrA | *S. oneidensis* MR-1 knocking out the gene *MtrA*, harbouring the empty plasmid PYYDT | Lab stock |
| OmcS/ΔMtrA | *S. oneidensis* MR-1 knocking out the gene *MtrA*, harbouring the plasmid $P_{tac}$-*OmcS*-PYYDT | This work |
| ΔMtrB | *S. oneidensis* MR-1 knocking out the gene *MtrB*, harbouring the empty plasmid PYYDT | Lab stock |
| OmcS/ΔMtrB | *S. oneidensis* MR-1 knocking out the gene *MtrB*, harbouring the plasmid $P_{tac}$-*OmcS*-PYYDT | This work |

| | | | |
|---|---|---|---|
| MKKGMKVSLS | VAAAALLMSA | PAAFAFHSGG | VAECEGCHTM |
| HNSLGGAVMN | SATAQFTTGP | MLLQGATQSS | SCLNCHQHAG |
| DTGPSSYHIS | TAEADMPAGT | APLQMTPGGD | FGWVKKTYTW |
| NVRGLNTSEG | ERKGHNIVAG | DYNYVADTTL | TTAPGGTYPA |
| NQLHCSSCHD | PHGKYRRFVD | GSIATTGLPI | KNSGSYQNSN |
| DPTAWGAVGA | YRILGGTGYQ | PKSLSGSYAF | ANQVPAAVAP |
| STYNRTEATT | QTRVAYGQGM | SEWCANCHTD | IHNSAYPTNL |
| RHPAGNGAKF | GATIAGLYNS | YKKSGDLTGT | QASAYLSLAP |
| FEEGTADYTV | LKGHAKIDDT | ALTGADATSN | VNCLSCHRAH |
| ASGFDSMTRF | NLAYEFTTIA | DASGNSIYGT | DPNTSSLQGR |
| SVNEMTAAYY | GRTADKFAPY | QRALCNKCHA | KD |

**Figure S1.** Mass spectrometry identification of OmcS peptides in the digested protein extracts. Peptide masses were queried against entries in the *Geobacter sulfurreducens* PCA database. The OmcS peptides highlighted in yellow were recovered.

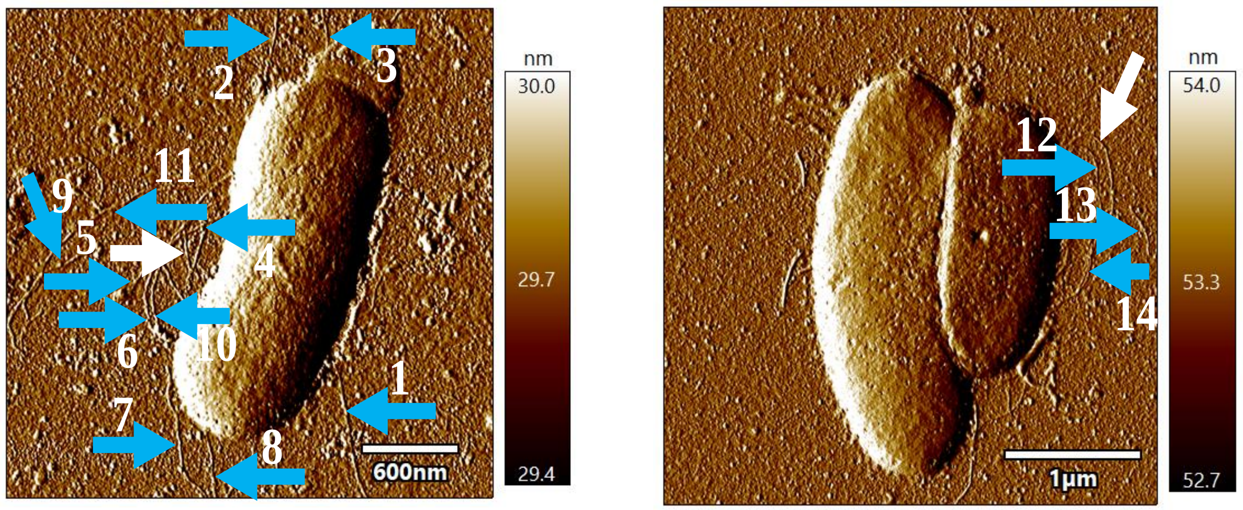


**Figure S2**. Illustration of relative abundance of OmcS filaments emanating from cells of strain OmcS. White arrow arrows point filaments with OmcS morphology (4 nm diameter, 20 nm longitudinal pitch) and are shown in more detail in the main text. Numbered blue arrows point to filaments without OmcS characteristics. Numbers on arrows refer to the numbered panels in figure S3 showing more detailed data on each filament.

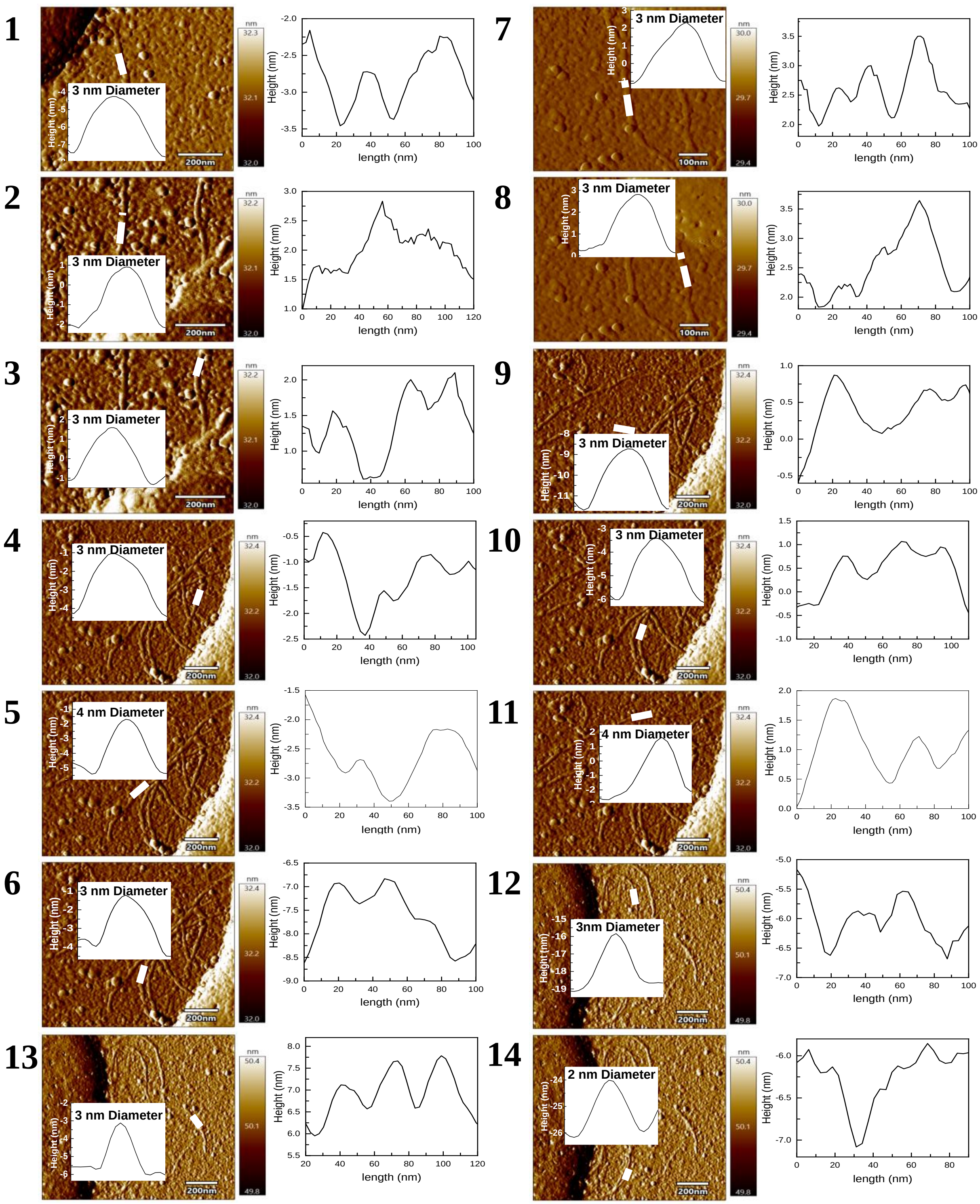

**Figure S3.** Morphology details of numbered filaments designated with a blue arrow in Figure S2.

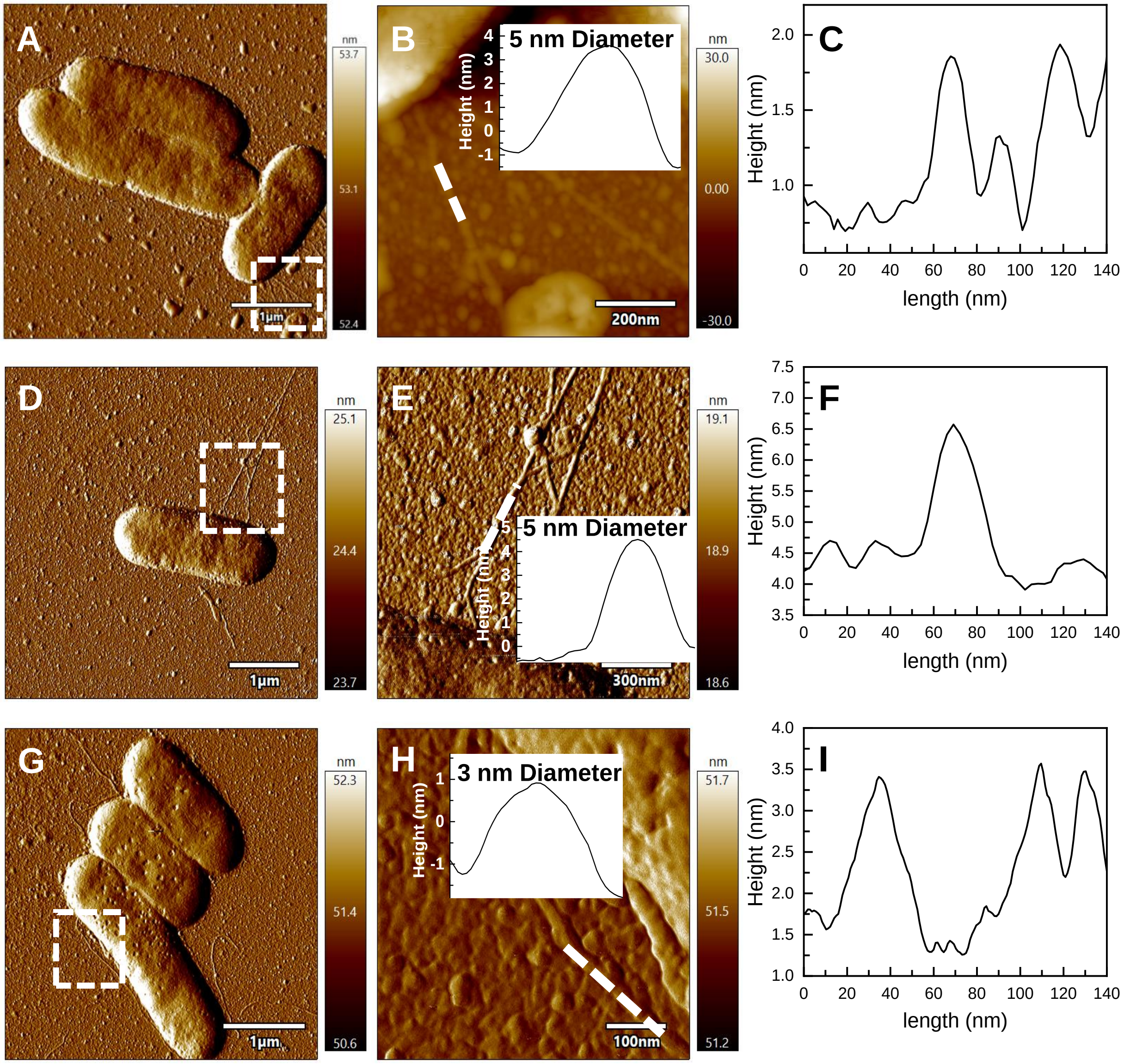


**Figure S4.** Examples of relative of typical filaments emanating from control strain MY.

### References


Aebersold, R., & Mann, M. (2003). Mass spectrometry-based proteomics. *Nature, 422*, 198-207.

Baron, D. B., LaBelle, E., Coursolle, D., Gralnick, J. A., & Bond, D. R. (2009). Electrochemical measurements of electron transfer kinetics by *Shewanella oneidensis* MR-1. *J. Biol. Chem., 284*, 28865-28873.

Bretschger, O., Obraztsova, A., Sturm, C. A., Chang, I. S., Gorby, Y. A., Reed, S. B., . . . Nealson, K. H. (2007). Current production and metal oxide reduction by *Shewanella oneidensis* MR-1 wild type and mutants. *Appl. Environ. Microbiol., 73*, 7003-7012.

Foster, L. J., & Mann, M. (2006). Protein identification and sequencing by mass spectrometry. In J. E. Celis (Ed.), *Cell Biology* (Vol. 4, pp. 363-369). London: Elsevier.

Grote, A., Hiller, K., Scheer, M., Münch, R., Nortemann, B., Hempel, D. C., & Jahn, D. (2005). JCat: a novel tool to adapt codon usage of a target gene to its potential expression host *Nucleic Acids Research, 33*, W526-W531.

Gstaiger, M., & Aebersold, R. (2009). Applying mass spectrometry-based proteomics to genetics, genomics and network biology. *Nature Reviews Genetics, 10*, 617-627.

Gundry, R. L., White, M. Y., Murray, C. I., Kane, L. A., Fu, Q., Stanley, B. A., & Van Eyk, J. E. (2010). Preparation of proteins and peptides for mass spectrometry analysis in a bottom-up proteomics workflow. In F. M. Ausubel (Ed.), *Current protocols in molecular biology* (pp. Chapter 10, Unit 10.25).

Heukeshoven, J., & Dernick, R. (1985). Simplified method for silver staining of proteins in polyacrylamide gels and the mechanism of silver staining. *Electrophoresis, 6*, 103-112.

Kanehisa, M., & Goto, S. (2000). KEGG: kyoto encyclopedia of genes and genomes. *Nucleic Acids Res., 28*(1), 27-30.

Liu, X., Walker, D. J. F., Nonnenmann, S., Sun, D., & Lovley, D. R. (2021). Direct observation of electrically conductive pili emanating from Geobacter sulfurreducens. *mBio, 12*, e02209-02221.

Ludwig, K. R., Schroll, M. M., & Hummon, A. B. (2018). Comparison of in-solution, fasp, and s-trap based digestion methods for bottom-up proteomic studies. . *Journal of Proteome Research*, *17*, 2480-2490.

Newman, D. K., & Saltikov, C. W. (2003). Genetic identification of a respiratory arsenate reductase. *Proc Natl Acad Sci U S A, 100*, 10983-10988.

Qian, X., Reguera, G., Mester, T., & Lovley, D. R. (2007). Evidence that OmcB and OmpB of *Geobacter sufurreducens* are outer membrane surface proteins. *FEMS Microb Lett*, *277*, 21-27.

Thomas, P. E., Ryan, D., & Levin, W. (1976). An improved staining procedure for the detection of the peroxidase activity of cytochrome P-450 on sodium dodecyl sulfate polyacrylamide gels. *Anal Biochem., 75*, 168-175.

Xie, Q., Lu, Y., Tang, L., Zeng, G., Yang, Z., Fan, C., . . . Atashgahi, S. (2020). The mechanism and application of bidirectional extracellular electron transport in the field of energy and environment. *Critical Reviews in Environmental Science and Technology, 51*, 1924-1969.

Yang, Y., Ding, Y., Hu, Y., Cao, B., Rice, S. A., Kjelleberg, S., & Song, H. (2015). Enhancing bidirectional electron transfer of Shewanella oneidensis by a synthetic flavin pathway. *ACS Synthetic Biology*, *4*, 815-823.
